# Ancient genomes reveal hybridisation between extinct short-faced bears and the extant spectacled bear (*Tremarctos ornatus*)

**DOI:** 10.1101/2021.02.05.429853

**Authors:** Alexander T Salis, Graham Gower, Blaine W. Schubert, Leopoldo H. Soibelzon, Holly Heiniger, Alfredo Prieto, Francisco J. Prevosti, Julie Meachen, Alan Cooper, Kieren J. Mitchell

**Author notes:** Corresponding author(s): A.T.S., and K.J.M.

## Abstract

Two genera and multiple species of short-faced bear from the Americas went extinct during or toward the end of the Pleistocene, and all belonged to the endemic New World subfamily Tremarctinae [1-7]. Two of these species were giants, growing in excess of 1,000 kg [6, 8, 9], but it remains uncertain how these extinct bears were related to the sole surviving short-faced bear: the spectacled bear (*Tremarctos ornatus*). Ancient mitochondrial DNA has recently suggested phylogenetic relationships among these lineages that conflict with interpretations based on morphology [1, 10-12]. However, widespread hybridisation and incomplete lineage sorting among extant bears mean that the mitochondrial phylogeny frequently does not reflect the true species tree [13, 14]. Here we present ancient nuclear genome sequences from representatives of the two extinct short-faced bear genera, *Arctotherium* and *Arctodus*. Our new data support a third hypothesis for the relationships among short-faced bears, which conflicts with existing mitochondrial and morphological data. Based on genome-wide D-statistics, we suggest that the extant spectacled bear derives substantial ancestry from Pleistocene hybridisation with an extinct short-faced bear lineage, resulting in a discordant phylogenetic signal between the mitochondrion and portions of the nuclear genome.

## Results and Discussion

The spectacled bear (*Tremarctos ornatus*) is the only extant species of short-faced bear (Tremarctinae), a once diverse subfamily endemic to the Americas. This subfamily also includes many species that became extinct during the Pleistocene, including the Florida cave bear (*Tremarctos floridanus*), two species of North American short-faced bears (*Arctodus* spp. [3, 4]), and as many as five species of South American short-faced bears (*Arctotherium* spp. [2, 6]), one of which (*Arctotherium wingei*) has recently been discovered as far north as the Yucatan of Mexico [5]. Notably, the genera *Arctodus* and *Arctotherium* both included giant (>1,000kg) forms [8, 9] — *Arctodus simus* and *Arctotherium angustidens*, respectively — and based on morphology it was hypothesised that these genera were closely related [1, 6, 10, 11]. However, recently published mitochondrial DNA data suggested that *Arctotherium* was most closely related to the extant spectacled bear, to the exclusion of North American *Arctodus* [12]. While this result supported the convergent evolution of giant bears in North and South America, the mitochondrial genome does not always reflect the true relationships among species [e.g. 15, 16-19]. Importantly, discordance between mitochondrial and nuclear loci has been previously noted in bears, and has been attributed to a combination of stochastic processes and the rapid evolution of bears [13], as well as hybridisation between species [13, 14, 20-25]. To further resolve the evolutionary history of short-faced bears, we applied ancient DNA techniques to retrieve and analyse whole genome data from both *Arctodus* and *Arctotherium*.

Ancient DNA (aDNA) was extracted and sequenced from three *Arctodus simus* specimens: one each from placer mines at Sixty Mile Creek (ACAD 438; Canadian Museum of Nature; CMN 42388) and Hester Creek (ACAD 344; Yukon Government; YG 76.4) in the Yukon Territory, Canada; and one from Natural Trap Cave in Wyoming, USA (ACAD 5177; University of Kansas; KU 31956). We also analysed one specimen of *Arctotherium* sp. from Cueva del Puma, Patagonia, Chile (ACAD 3599; complete right femur, no. 32104, Centro de Estudios del Hombre Austral, Instituto de la Patagonia, Universidad de Magallanes). The *Arctotherium* specimen was previously dated to 12,105 ± 175 cal yBP (Ua-21033) [26], while two of the *Arctodus* specimens have been dated: ACAD 438 at 47,621 ± 984 cal yBP (TO-2699) [27] and ACAD 5177 at 24,300 ± 208 cal yBP (OxA-37990) (Table S1). The *Arctotherium* specimen has yielded mitochondrial aDNA in a previous studies [12], however, here we shotgun sequenced this specimen, along with the three *A. simus* specimens, at much greater depth in order to reconstruct nuclear genome sequences. Mapping our new sequencing data from these specimens to the giant panda (*Ailuropoda melanoleuca*) reference genome (LATN01) yielded average depths of coverage between 0.12x to 5.9x for the *A. simus* specimens and 3.9x for the *Arctotherium* specimen (Table S3). We compared these new genomic data to previously published genomes from all extant species of bear (Table S2): spectacled bear, giant panda, brown bear (*Ursus arctos*), American black bear (*U. americanus*), Asian black bear (*U. thibetanus*), polar bear (*U. maritimus*), sloth bear (*U. ursinus*), and sun bear (*U. malayanus*).

Phylogenetic analyses on a concatenated dataset of genome-wide SNPs revealed relationships within Ursinae that were consistent with previous genomic studies: *U. americanus, U. maritimus*, and *U. arctos* formed a monophyletic clade sister to a clade consisting of *U. thibetanus, U. malayanus*, and *U. ursinus* [13, 14]. In contrast, within short-faced bears (Tremarctinae) we recovered strong support for a close relationship between the spectacled bear and the North American short-faced bear (*Arctodus simus*) to the exclusion of the South American *Arctotherium* (Figure 1A, Figure S2). This result conflicts with the mitochondrial tree, which instead supports a clade comprising *Arctotherium* and *Tremarctos ornatus* to the exclusion of *Arctodus simus* [12] (Figure 1B). As the radiation of bears is thought to have occurred rapidly during the Miocene - Pliocene transition, it is possible that this discordance could be explained by incomplete lineage sorting (ILS) [28], a process whereby pre-existing genetic variation in an ancestral species is randomly inherited and fixed in descendant species [29, 30]. Alternatively, given the observed propensity of bears for hybridisation [e.g. 13, 14, 20-22, 25, 31], mitochondrial/nuclear discordance within short-faced bears may instead result from gene flow between *Tremarctos* and either *Arctodus* or *Arctotherium*.

**Figure 1:**
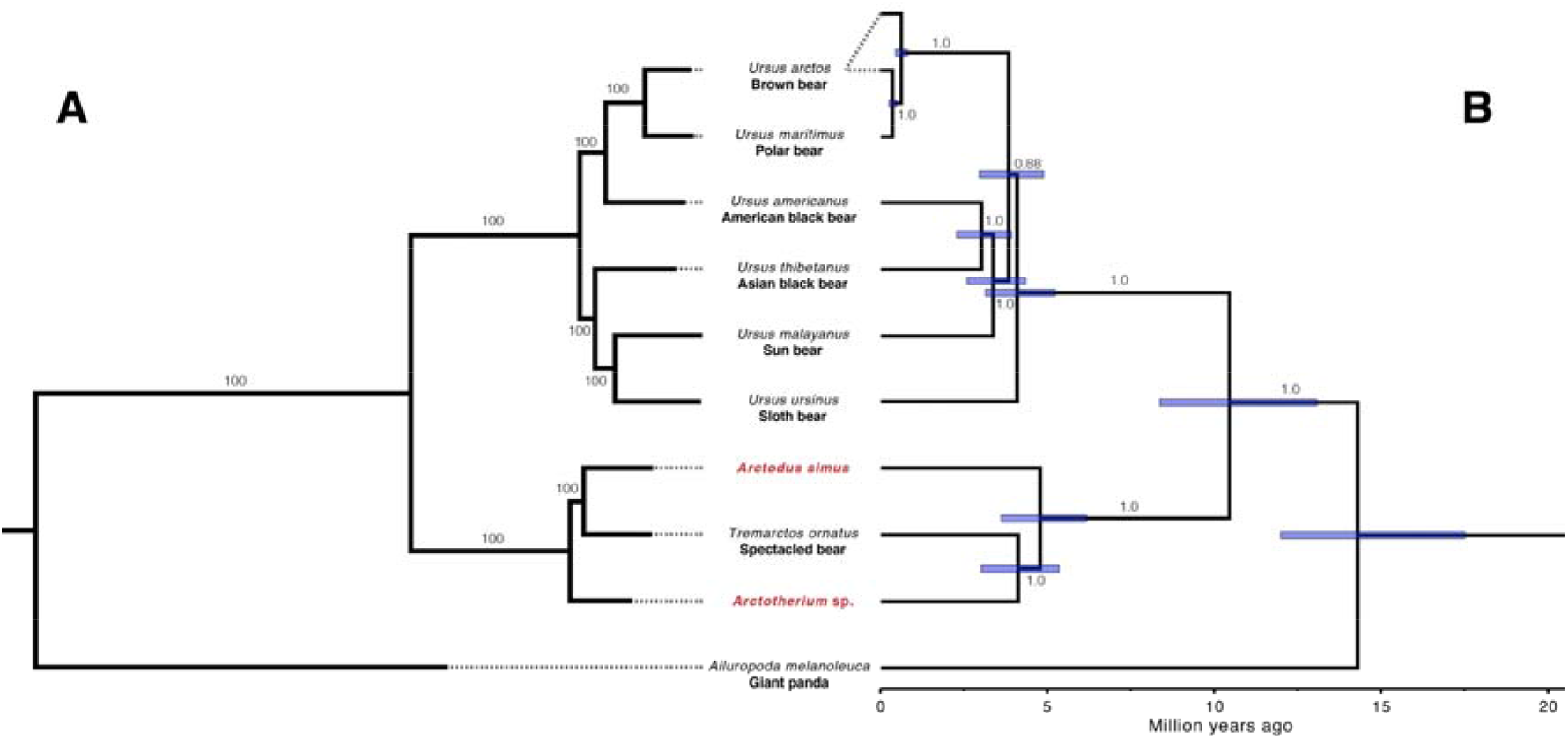
Phylogenetic relationships among ursids **A**. Maximum likelihood tree based on nuclear SNPs constructed in RAxML. Branch labels represent bootstrap support percentages. For RAxML tree with all individuals analysed see Figure S2. **B**. Bayesian phylogeny based on full mitochondrial genomes adapted from Mitchell, et al. [12]. Blue bars represent 95% highest posterior density interval on node ages. Branch labels represent BEAST posterior support values.

To test for potential phylogenetic discordance across our short-faced bear genomes, we constructed phylogenetic trees from 500 kb non-overlapping windows (n = 2622) across the 85 largest autosomal scaffolds of the giant panda reference genome (LATN01). Trees created from roughly 70% of windows agreed with the results from our genome-wide concatenated dataset (Topology 1; *i*.*e. Tremarctos* + *Arctodus*; Figure 2B & S3). However, approximately 30% of windows instead supported the mitochondrial tree topology (Topology 2; *i*.*e. Tremarctos* + *Arctotherium*; Figure 2B & S3), while the third possible topology where the two extinct genera form a clade *— Arctodus* + *Arctotherium —* was rejected for over 95% of windows. The frequencies of the three possible tree topologies are difficult to explain as a result of ILS, which we would expect to result in a more even representation of the two “minority” topologies (*i*.*e*. Topologies 2 and 3). Our results therefore suggest that introgression may be the most likely explanation for the observed phylogenetic discordance. Consequently, we calculated D-statistics [32, 33] using our concatenated genome-wide SNPs in order to identify signals of hybridisation between the bear species in our dataset.

**Figure 2:**
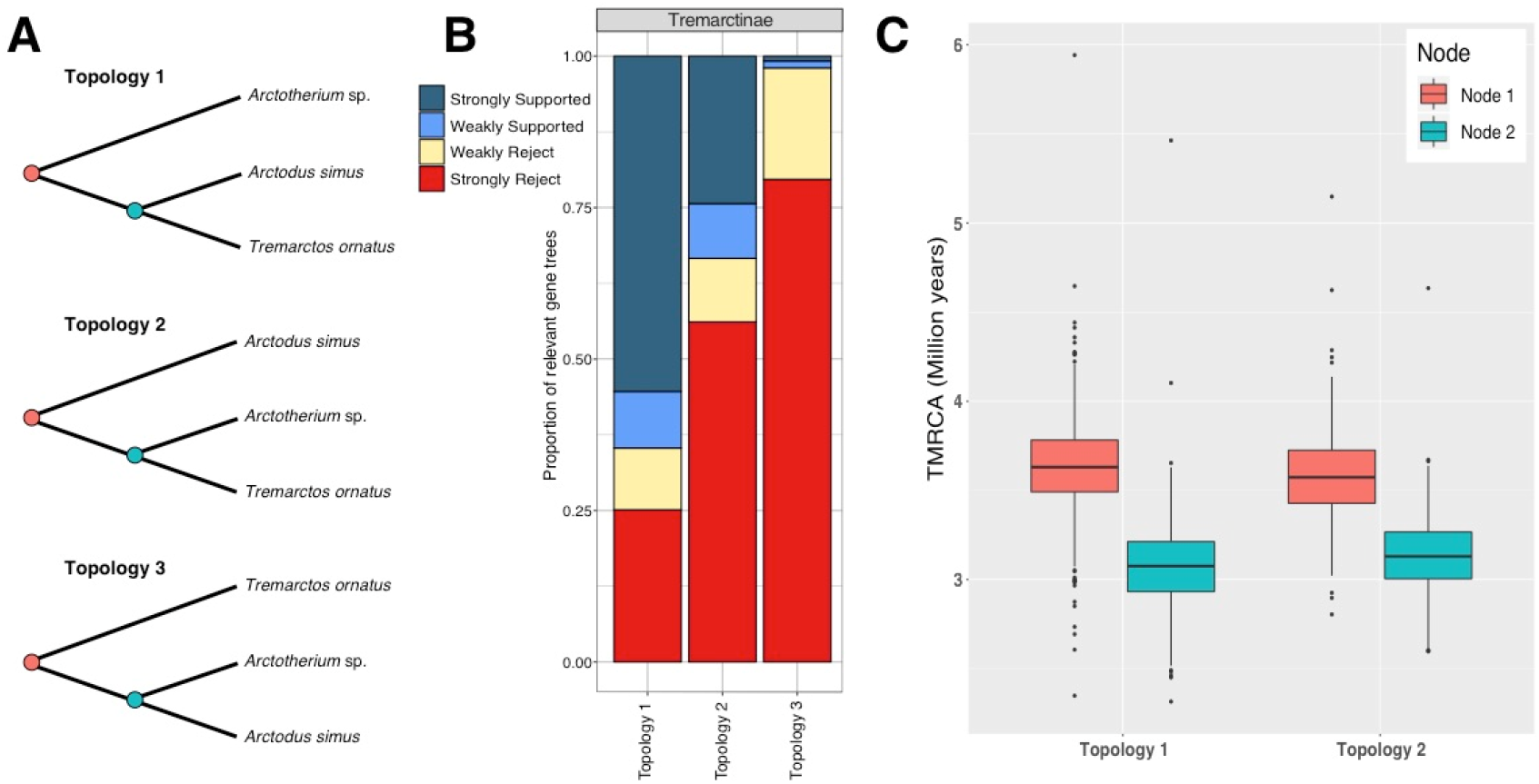
**A**. The three possible short-faced bear (Tremarctinae) tree topologies. **B**. Discordance visualisation using DiscoVista from 2622 500 kb genomic fragments. The x-axis represents topologies tested and the y-axis the proportion of fragments that support the topology, with >80% bootstrap support used to define strong support. For more comprehensive tests of phylogenetic placements see Figure S3. **C**. Divergence time estimates (TMRCA) of *Tremarctos, Arctodus, and Arctotherium* (Node 2) and for sister species (Node 1) of the two most common short-faced bear topologies.

Consistent with previous studies [i.e. 14], our D-statistics revealed compelling evidence for hybridisation between: Asian black bears and all North American ursine bears (including the polar bear); sun bears and North American ursine bears; and Asian black bear and sun bear (Table S4). In contrast, we did not obtain any significantly non-zero values for D-statistics calculated using our two extinct short-faced bear genomes, any member of Ursinae, and the panda outgroup (Table 1). This result suggests that no gene flow occurred between *Arctodus* or *Arctotherium* and the ancestors of any modern ursine bear, and also demonstrates a lack of any discernible reference bias in the ancient genomic data (which would result in asymmetrical allele sharing with the reference). Thus, it appears *Arctodus* and *Arctotherium* did not hybridise with brown and black bears in the Americas during the late Pleistocene, even though the distribution of *Arctodus* overlapped with both ursines, and *Arctotherium* may have encountered them in Mexico or Central America [5]

Contrary to previous studies, our D-statistics revealed signals consistent with gene flow between the spectacled bear and members of Ursinae (Table 1 & S5), suggesting the possibility that *Tremarctos* hybridised with ancestors of either the brown bear or American black bear during the Pleistocene. This signal is surprising given the deep divergence between ursine and short-faced bears, having split approximately 10 million years ago (mya) [12, 14, 28]. However, in support of this hypothesis, offspring between spectacled bear and Asiatic black bear have resulted from hybridisation in zoos, although whether these hybrids were fertile remains unknown [34]. Importantly, members of *Tremarctos* and the ancestors of modern American black bears had overlapping distributions throughout the Pleistocene in North America [4, 10], meaning that hybridisation may have occurred when the two lineages were less divergent and reproductive barriers had had less time to evolve.

**Table 1:**
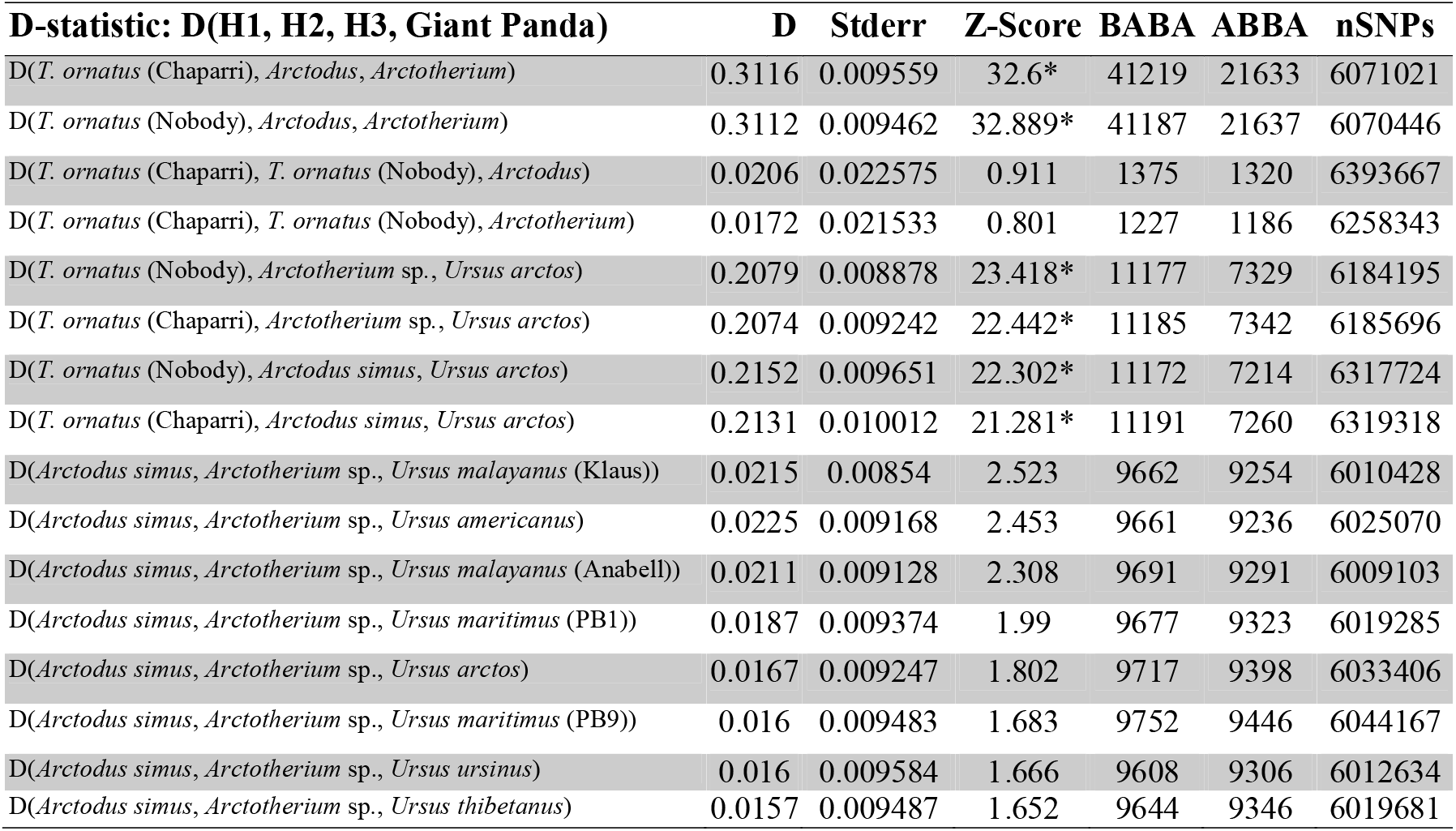
D-statistics for short-faced bears (Tremarctinae). D-statistics (D), standard error, and Z-Score (significant if > |3|) are displayed, with ABBA-BABA counts and the number of SNPs considered in the analysis. It is clear that there is an excess of allele sharing between the spectacled bears (*T. ornatus*) and *Arctotherium*. However, neither of the spectacled bear individuals show elevated D-statistics in relation to each other meaning gene flow likely occurred in the ancestor of both individuals, or they carry similar proportions of hybridised DNA. *Significantly positive d-statistic, denote deviations from a typical bifurcating tree with H1 and H3 being closer than expected

In addition to evidence for hybridisation between *Tremarctos* and ursine bears, we also recovered convincing evidence for hybridisation between *Arctotherium* and *Tremarctos* (Table 1). These results are consistent with a model where the divergence between *Arctodus* and *Tremarctos* occurred in North America after the ancestors of *Arctotherium* dispersed southwards into South America, with subsequent hybridisation between *Tremarctos* and *Arctotherium*. This interpretation is supported by the presence of *Arctodus* and *Tremarctos* (and absence of *Arctotherium*) in the late Pliocene fossil record of North America [3, 4, 7, 10]. The fossil record further suggests that contact between *Tremarctos* and *Arctotherium* occurred during the late Pleistocene, when representatives of *Arctotherium* were distributed as far north as the Yucatan of Mexico [5], providing an opportunity for hybridisation.

If the ancestors of the spectacled bear hybridised with *Arctotherium* somewhere in the American mid-latitudes during the migration of *Tremarctos* into South America, then signals of gene flow between members of these two genera could date to the latest Pleistocene or earliest Holocene, when spectacled bears are thought to have migrated into South America [6, 35, 36]. To test this hypothesis, we estimated divergence times among the three short-faced bear lineages for all 500 kb windows from the largest 40 scaffolds corresponding to either Topology 1 (n = 980) or Topology 2 (n = 413) and summarised the results (Figure 2c). The age of the most recent common ancestor (TMRCA) of *Tremarctos, Arctodus, and Arctotherium* was similar irrespective of topology (Topology 1: 3.6 mya; Topology 2: 3.6 mya), as was the subsequent divergence between the remaining two lineages (Topology 1: 3.1 mya; Topology 2: 3.1 mya). Assuming that members of *Tremarctos* migrated southward no earlier than the latest Pleistocene, our results superficially appear to be incompatible with late Pleistocene/Holocene hybridisation between *Tremarctos* and *Arctotherium*. The fossil record suggests two ways these observations may be explained.

Late Pleistocene fossil data indicate that the ancestors of the spectacled bear are likely to have encountered *Arctotherium* individuals from Mexico, Central America, and/or northern South America, which were comparable in size and diet to the spectacled bear [1, 5, 37] and which may have represented a different *Arctotherium* species from the Chilean specimen sequenced in the present study [1, 6, 12, 26]. Indeed, throughout the Pleistocene a number of *Arctotherium* species have been described across South and Central America, with putative species ranging from gigantic in the early-mid Pleistocene to relatively small in the late Pleistocene [1, 2, 6, 9]. If the ancestors of our sampled Patagonian *Arctotherium* specimen diverged from those of more northerly *Arctotherium* species during the Pliocene or early Pleistocene, then our molecular dating results remain consistent with hybridisation being the primary driver of phylogenetic discordance in our genomic data. Alternatively, hybridisation between *Tremarctos* and *Arctotherium* could have occurred in Central America during the Pleistocene. *Tremarctos* and *Arctotherium* have both been recorded in Central American cave deposits [5, 38], however, the extent of occupation by both genera in the region is unknown, and conceivably Central America represents a contact zone between the genera throughout the Pleistocene where hybridisation may have occurred.

An alternative interpretation of our phylogenetic results is that Topology 2 (*Tremarctos* + *Arctotherium*), which is supported by the mitochondrion and ∼30% of our nuclear genome windows, is the pre-hybridisation tree. Recently, Li, et al. [39] suggested that under scenarios involving substantial gene flow the predominant phylogenetic signal across the genome may not reflect the pre-hybridisation tree. If this were the case for short-faced bears, the majority of support for Topology 1 would actually result from extensive hybridisation between *Arctodus* and *Tremarctos* in North America. Li, et al. [39] contend that the phylogenetic signal of the pre-hybridisation tree may be enriched in regions of low recombination, especially on the X-chromosome. In order to test this hypothesis, we identified panda scaffolds corresponding to the ∼40 Mb recombination cold-spot on the X-chromosome highlighted by Li, et al. [39] and produced phylogenetic trees for each 500 kb window along this region (Figure S4). Interestingly, the majority of these fragments supported Topology 2 (*Tremarctos* + *Arctotherium*), the same topology as the mitochondrial phylogeny but contrasting with the majority of autosomal scaffolds.

Unlike felids [e.g. 40, 41, 42], a high-quality reference assembly and linkage map does not exist for any bear species, meaning scaffolds pertaining to high and low recombination areas of the genome could not be identified. Unfortunately, this currently makes it impossible to further explore the possibility that Topology 2 (*Tremarctos* + *Arctotherium*) may reflect the pre-hybridisation short-faced bear tree, rather than Topology 1 (*Tremarctos* + *Arctodus*). In the absence of a linkage map, sequencing aDNA from either the extinct *Tremarctos floridanus* or more northerly *Arctotherium* populations will be key to further resolving the evolutionary history of short-faced bears, though this will be challenging given that the core range of these species lies in the lower-latitudes where aDNA preservation is less reliable. For now we conclude that the weight of evidence supports a closer relationship between the spectacled bear and the extinct short-faced bears from North America (*Arctodus*) rather than South America (*Arctotherium*). In any case, our genomic data imply extensive hybridisation occurred between the spectacled bear and one of the extinct short-faced bear lineages. These results contribute to the growing consensus that hybridisation is widespread among carnivoran groups generally [13, 14, 39, 43].

## Acknowledgements

We would like to thank the following institutions for allowing access to specimens: Canadian Museum of Nature, University of Kansas Natural History Museum, Yukon Government, Centro de Estudios del Hombre Austral, Instituto de la Patagonia, Universidad de Magallanes. In addition, we are grateful to the following individuals who helped in the collection and identification of specimens and/or provided laboratory support: Grant Zazula (Yukon Territorial Government, Palaeontology Program, Canada), Fabiana Martin (Universidad de Magallanes, Chile), Jeremy Austin (University of Adelaide, Australia) and Sarah Bray (University of Adelaide, Australia). We would like to thank the Wyoming BLM and permit number PA-13-WY-207. This research was funded by an Australian Research Council Laureate Fellowship awarded to AC (FL140100260), U.S. National Science Foundation grant (EAR/SGP#1425059) awarded to JM and AC, and Agencia Nacional de Promoción Científica y Técnica’ (ANPCyT, Argentina) (PICT 2015–966) awarded to FJP.

## Author Contributions

Conceptualization, A.T.S., A.C. and K.J.M.; Methodology, A.T.S., G.G., A.C., and K.J.M.; Investigation, A.T.S., H.H., and K.J.M.; Writing – Original Draft, A.T.S. and K.J.M.; Writing – Review & Editing, G.G., B.W.S., L.H.S., H.H., A.P., F.J.P., J.M., and A.C.; Funding Acquisition, F.J.P., J.M., and A.C.; Resources, J.M., A.P., and F.J.P.; Supervision, A.C., and K.J.M.

## Materials and Methods

### Sampling

Analyses were performed on three bone samples identified as *Arctodus simus* and one sample identified as *Arctotherium* sp. (Table S1). The *Arctotherium* specimen ACAD 3599, had previously be radiocarbon dated, as well as one of the *A. simus* specimens (ACAD 438), a further *A. simus* specimen was radiocarbon dated at the Oxford Radiocarbon Accelerator Unit of the University of Oxford. All radiocarbon dates were calibrated with the either the IntCal13 curve [44] or the SHCal13 curve [45] using OxCal 4.4 [46] (Table S1).

### Sample preparation and extraction

All pre-PCR steps (*i*.*e*., extraction, library preparation) were conducted in purpose-built ancient DNA clean-room facilities at the University of Adelaide’s Australian Centre for Ancient DNA (ACAD). Potential surface contamination on each sample was reduced by UV irradiation for 15 min each side, followed by abrasion of the exterior surface (c. 1 mm) using a Dremel tool and a disposable carborundum disk. The sample was then pulverised using a metallic mallet. Approximately 100 mg of powder was extracted using an in-house silica-based extraction protocol adapted from Dabney, et al. [47] optimised for the recovery of small fragments. the powder was digested first in 1 mL 0.5 M EDTA for 60 min, followed by an overnight incubation in 970 μL fresh 0.5 M EDTA and 30 μL proteinase K (20 mg/ml) at 55°C. The samples were centrifuged and the supernatant mixed with 13 mL of a modified PB buffer (12.6 mL PB buffer (Qiagen), 6.5 μL Tween-20, and 390 μL of 3M Sodium Acetate) and bound to silicon dioxide particles, which were then washed two times with 80% ethanol. The DNA was eluted from silica particles with 100 μL TE buffer.

### Library preparation

Double-stranded Illumina libraries were constructed following the protocol of Meyer, et al. [48] from 25 μL of DNA extract. In addition, all samples underwent partial uracil-DNA glycosylase (UDG) treatment [49] to restrict cytosine deamination, characteristic of ancient DNA, to terminal nucleotides. A short round of PCR using PCR primers complementary to the library adapter sequences was performed to increase the total amount of DNA and add full-length Illumina sequencing adapters. Cycle number was determined via rtPCR and each library split into 8 separate PCR reactions to minimise PCR bias and maintain library complexity. Each PCR of 25 μL contained 1× HiFi buffer, 2.5 mM MgSO4, 1 mM dNTPs,each primer, 0.1 U Platinum Taq Hi-Fi polymerase and 3 μL DNA. The cycling conditions were 94 °C for 6 min, 8–10 cycles of 94 °C for 30 s, 60 °C for 30 s, and 72 °C for 40 s, followed by 72 °C for 10 min. Following PCR, replicates were pooled and purified using AxyPrep™ magnetic beads, eluted in 30 μL H2O quantified on TapeStation (Agilent Technologies).

### Sequencing

Libraries were initially pooled and sequenced on an Illumina NextSeq using 2 x 75 bp PE (150 cycle) High Output chemistry. For deeper sequencing, libraries were diluted to 1.5 nM and each was run on one lane of an Illumina HiSeq X Ten using 2 x 150 bp PE (300 cycle) chemistry, except for ACAD 438 which was run on two lanes of an Illumina HiSeq X Ten.

### Data processing

Demultiplexed sequencing reads were processed through Paleomix v1.2.12 [50]. Within Paleomix, raw reads were filtered, adapter sequences removed, and pair-end reads merged using ADAPTER REMOVAL v2.1.7 [51], trimming low quality bases (<Phred20 -- minquality 4) and discarding merged reads shorter than 25 bp (--minlength 25). Read quality was visualised before and after adapter trimming using fastQC v0.11.5 (http://www.bioinformatics.babraham.ac.uk/projects/fastqc/) to ensure efficient adapter removal. Reads were mapped to the Panda ASM200744v1 genome [52] with BWA v0.7.15 using the mem algorithm [53]. Reads with mapping Phred scores less than 25 were removed using SAMtools 1.5 [54] and PCR duplicates were removed using “paleomix rmdup_collapsed” and MARKDUPLICATES from the Picard package (http://broadinstitute.github.io/picard/). Indel realignment was performed using GATK [55] and damage profiles assessed using MapDamage v2.0.8 [56] (Figure S1).

Sequencing reads were downloaded from the European Nucleotide Archive for all extant bear species (Table S2) [14, 21, 24, 57, 58] and processed using the same pipeline as for the ancient samples.

### Phylogenetic analysis

Indexed VCF files were created for each BAM file using mpileup, part of the SAMtools package v0.1.19 [54], and the call and index functions as a part of the BCFtools package v0.1.19. Parallel v2010622 [59] was used to process each BAM file in parallel. BCFtools was then used to filter SNPs within 3 bp of an indel (--SnpGap 3). The 85 largest scaffolds of the Panda reference genome were renamed as chromosomes (chr1–85) in each VCF file using BCFtools annotate. Biallelic variants in VCF files were converted to random pseudohaploid variants in eigenstrat format for the 85 largest scaffolds using vcf2eig (part of eig-utils; https://github.com/grahamgower/eig-utils) including monomorphic (-m) and singleton (-s) sites, and excluding transitions (-t). Eigenstrat formatted files were then converted to PHYLIP files using eig2phylip (part of eig-utils; https://github.com/grahamgower/eig-utils). A supermatrix tree was then created in RAxML v8.2.4 [60] using the rapid bootstrapping algorithm (-f a) and using the GTRCAT model of substitution with ascertainment correction (-m ASC_GTRCAT) with 100 bootstrap replicates (-#100) and using the Felsenstein ascertainment correction (--asc-corr=felsenstein) based on the number of invariant sites (calculated from the total ungapped length of the largest 85 scaffolds of the Panda reference genome minus the length of the alignment).

### Discordance analysis using DiscoVista

The eigenstrat files were broken down into non-overlapping 500kb sliding windows using eigreduce (part of eig-utils; https://github.com/grahamgower/eig-utils). We only used the higher coverage A. simus sample (ACAD 344) in these analyses. For each window a PHYLIP file and tree were created as described above. The frequency and support of different tree topologies was then summarised and visualised using DiscoVista [61], using bootstrap values of 80 as the cutoff for strong support. Topologies tested included: 1) the inclusion of *Arctotherium* with ursine bears; 2) the inclusion of *Arctodus* with ursine bears; 3) the inclusion of *Tremarctos* with ursine bears; 4) any combination of tremarctine bears included with ursine bears; 5) the monophyly of Tremarctinae; 6) monophyly of *Tremarctos* and *Arctodus*; 7) monophyly of *Tremarctos* and *Arctotherium*; and 8) monophyly of *Arctotherium* and *Arctodus*.

### D-statistics

To test for signals of gene flow within Tremarctinae and between tremarctine and ursine lineages we used D-statistics as implemented by Admixtools [62] in admixr [63]. We only used the higher coverage *A. simus* sample (ACAD 344) in this analysis. The giant panda was used as outgroup and block jack-knife procedure used to test for significant departures from zero (|Z|>3). D-statistics within Tremarctinae were calculated in the form D(*Arctodus, Tremarctos, Arctotherium*, panda) and for detecting gene flow between Tremarctinae and Ursinae in the form D(U1, U2, T1, panda), where T1 is any short-faced bear and U1 and U2 any ursine individual. D-statistics were also performed to detect gene flow within Ursinae (as per Kumar, et al. 2017), using either the giant panda or spectacled bear as outgroup. To account for the possibility of a reference bias in ancient samples, within Tremarctinae D-statistics were recalculated using the Asiatic black bear as outgroup (Table S5).

### Molecular dating

Divergence times were estimated for each 500kb fragment from the discordance analysis using MCMCtree, part of the PAML package v4.8a [64], using the topology from the ML tree produced in the discordance analysis as the input tree. Four calibrations were used to calibrate the phylogeny:

1. The crown-age of Ursidae (*i*.*e*. the divergence of the giant panda lineage) was constrained to between 11.6 and 23 million years ago (mya) based on the presence of *Kretzoiarctos* [65], a putative ailuropodine, in the middle Miocene and the assumption that early Miocene *Ursavus* representatives are likely ancestral to modern ursids [10].
2. The divergence of Tremarctinae and Ursinae was constrained to between 7 and 13 mya based on the presence of putative early tremarctine bears (*e*.*g. Plionarctos*) in the Late Miocene/Early Pliocene [66].
3. The common ancestor of all sampled ursine bears was constrained to between 4.3 and 6 mya based on the occurrence of *Ursus minimus* [14, 67].
4. The divergence of polar and brown bears was constrained to between 0.48 and 1.1 mya based on previous nuclear estimates [21, 24, 25].

The JC +G substitution model with 5 discrete gamma categories was used with autocorrelated-rates, also known as the geometric Brownian diffusion clock model. Uniform priors for node ages using the birth-death (BD) process were used [λ_BD_ = 1 (birth-rate), μ_BD_ = 1 (death-rate), and ρ_BD_ = 0.1 (sampling fraction for extant species)]. A gamma-Dirichlet distribution was used for the prior on rate with an α shape parameter of 2 (diffuse prior). The 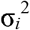 prior was defined as a diffuse gamma-Dirichlet distribution (2,2). MCMC tree runs were performed with a burn-in of 10000, and a sample size of 10000, sampling every ten iterations. Median node ages were then averaged for each tree topology.

### Low-recombining region of X-chromosome

Scaffolds of the panda ASM200744v1 reference genome [52] corresponding to low recombination regions of the X chromosome were identified by mapping all scaffolds to the recombination cold-spot of the X chromosome of the domestic cat (FelCat5) using minimap2. Default parameters were used, meaning the alignment lacked base-level precision (to account to phylogenetic distance between giant panda and the domestic cat). Only scaffolds larger than 500kb and with greater than 100 kb of segments mapping to the low recombining region of the domestic cat X-chromosome were retained, resulting in 15 scaffolds linked to the low recombination region of the X-chromosome. A maximum-likelihood phylogenetic tree and gene-tree discordance analysis were performed on these 15 scaffolds as described above for the genome-wide dataset.

## Notes

### Competing Interest Statement

The authors have declared no competing interest.

### Summary of Updates

Corrections to an incorrect species reference in the citation of published literature and mistakes in the reporting of sequencing and mapping statistics in the supplementary information.

