## Supplementary Information for "Ancient genomes reveal hybridisation between extinct short-faced bears and the extant spectacled bear (*Tremarctos ornatus*)"

Supplementary Figures S1-S4

Supplementary Tables S1-S5

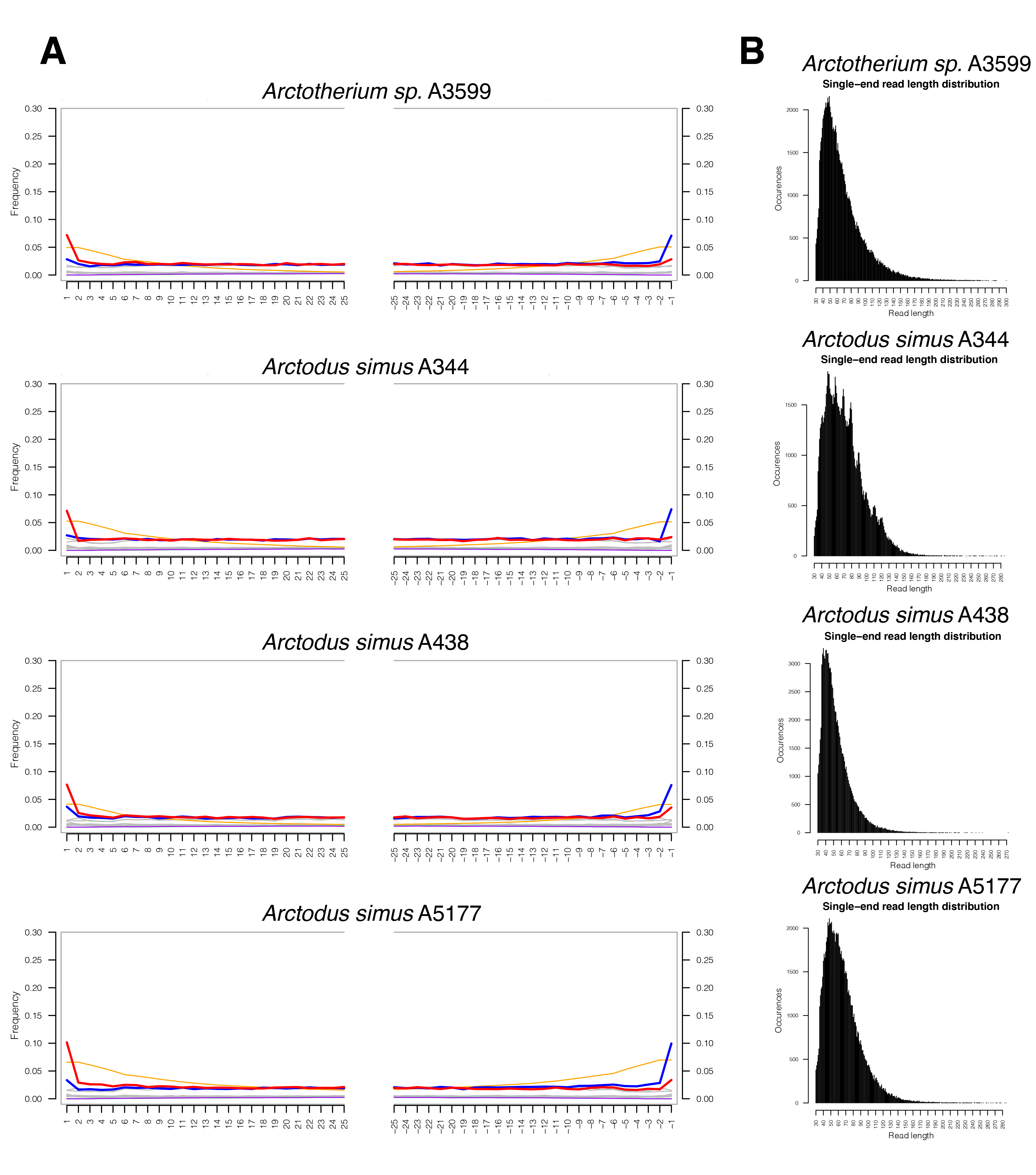

­

**Figure S1:** Authentication of ancient genomic data from the four extinct short-faced bears specimens. **A)** Cytosine deamination patterns where the x-axis represents the number of bases from the 5’ or 3’ end of a DNA fragment. The red lines represent cytosine (C) to thymine (T) transitions and blue lines guanine (G) to adenine (A) transitions compared to the giant panda reference genome. All samples show and accumulation of C-T transitions on the first base of DNA fragments, characteristic of aDNA that has undergone partial UDG treatment. The increase in G-A at terminal 3’ base is an artefact of the double-stranded DNA library procedure and actually represents C-T transitions. **B)** Fragment length distributions of the length of reads mapping to the giant panda reference from each sample. The minimum read length used for mapping was 25 bp, resulting in a hard cut-off at this length. All samples show an abundance of small fragments (all centred around ~50 bp), characteristic of aDNA.

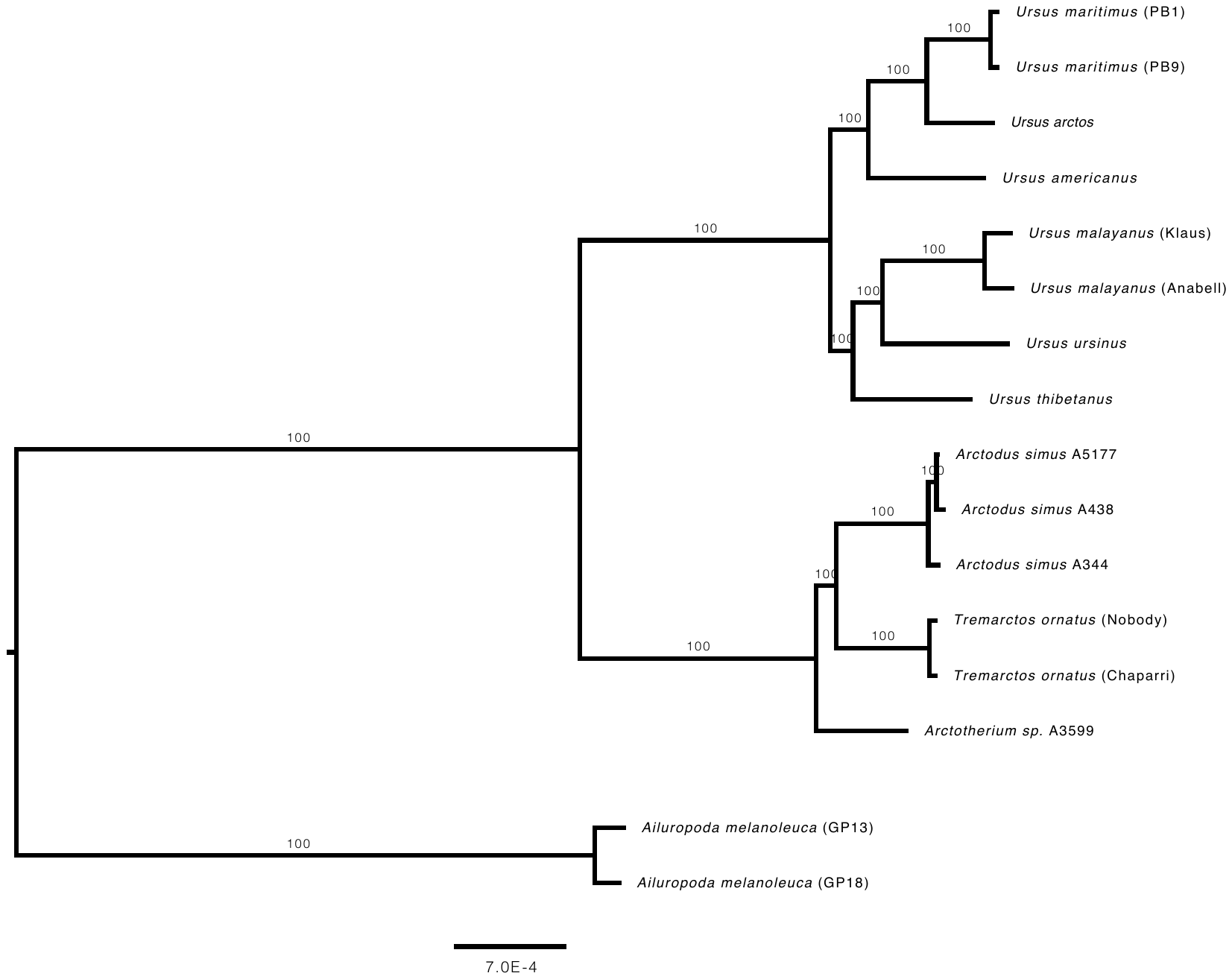

**Figure S2:** Maximum likelihood tree based on nuclear SNPs constructed in RAxML. Branch labels represent bootstrap support percentages.

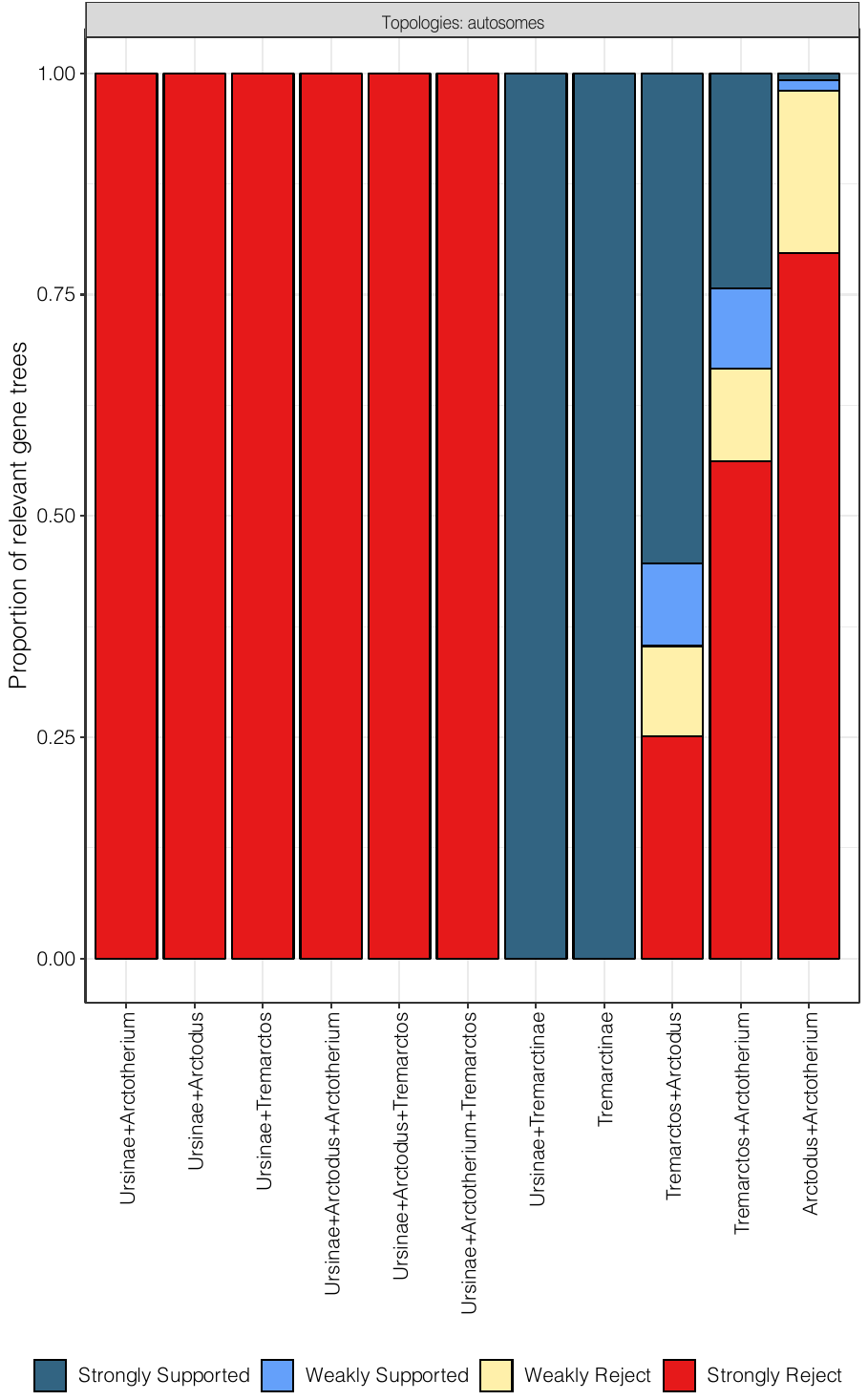

**Figure S3:** Discordance visualisation using DiscoVista from 2622 500 kb autosomal genomic fragments for all tested topologies involving tremarctine bears, with >80 bootstrap support used to define strong support. The x-axis represents topologies tested and the y-axis the proportion of fragments that support the topology.

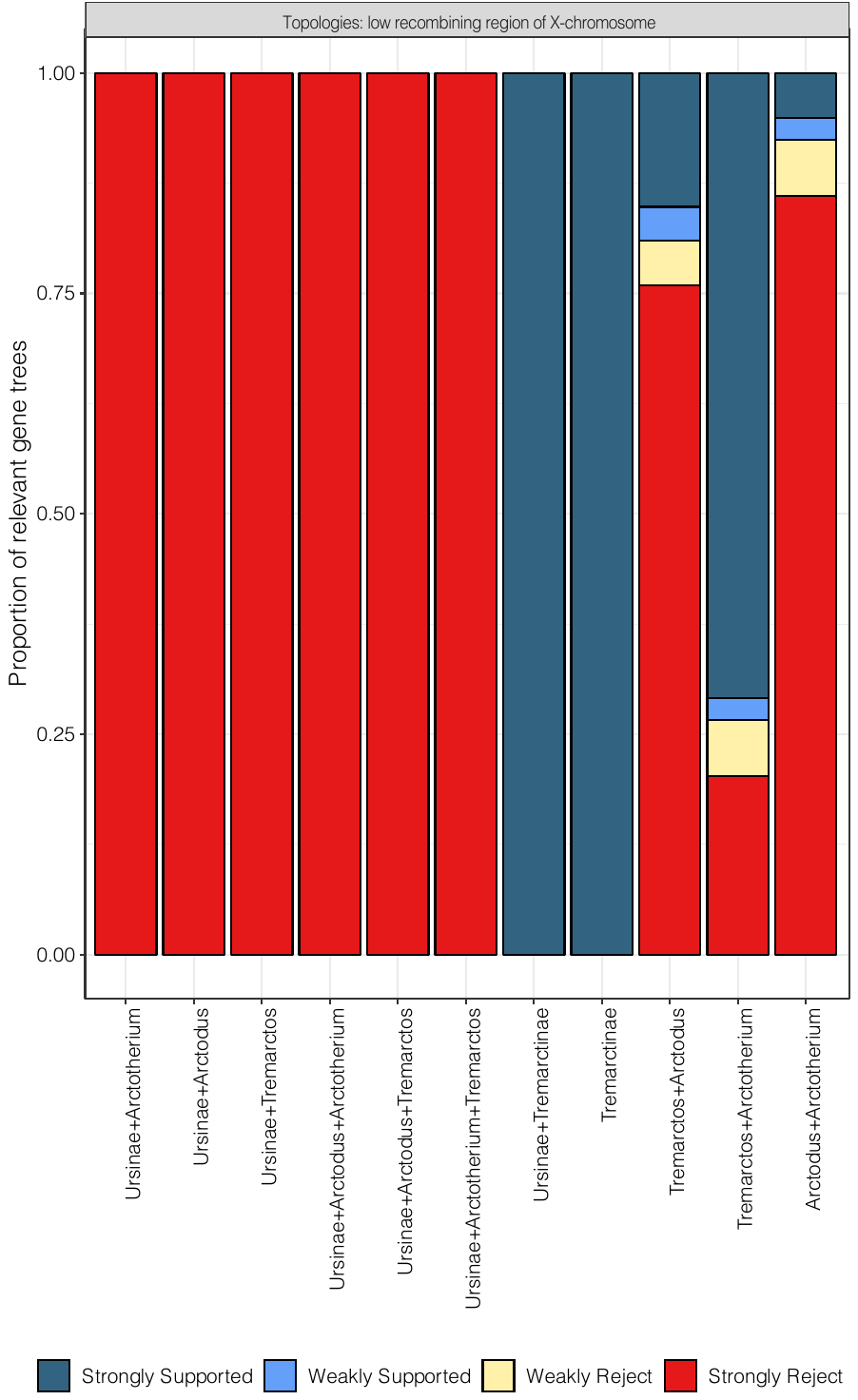

**Figure S4:** Discordance visualisation using DiscoVista from 80 500 kb genomic fragments pertaining to the ~40 Mb recombination cold-spot on the X chromosome, with >80 bootstrap support used to define strong support. The x-axis represents topologies tested and the y-axis the proportion of fragments that support the topology.

**Table S1:** Radiocarbon dates of samples.

| Sample Name | Radiocarbon date | Radiocarbon reference | Calibrated  date | OxCal calibration curve | Study |
| --- | --- | --- | --- | --- | --- |
| ACAD 438 | 44240 ± 930 | TO-2699 | 47621 ± 984 | IntCal13 | Harington, et al. [1] |
| ACAD 3599 | 10345 ± 75 | Ua-21033 | 12105 ± 175 | SHCal13 | Martin, et al. [2] |
| ACAD 5177 | 20220 ± 150 | OxA-37990 | 24300 ± 208 | IntCal13 | this study |

| Binomial Name | Common Name | Sample Name | Location | EBI Sample Accession | EBI Read Accession | Study |
| --- | --- | --- | --- | --- | --- | --- |
| **Arctodus simus* | North American short-faced bear | ACAD 344/ YG 76.4 | Hester Creek, Yukon | NA | NA | this study |
| **Arctodus simus* | North American short-faced bear | ACAD 438/ CMN 42388 | Sixty Mile Creek, Yukon | NA | NA | this study |
| **Arctodus simus* | North American short-faced bear | ACAD 5177/ KU 31956 | Natural Trap Cave, Wyoming | NA | NA | this study |
| **Arctotherium sp.* | South American short-faced bear | ACAD 3599/ no. 32104 | Cueva del Puma, Patagonia | NA | NA | this study |
| *Tremarctos ornatus* | Spectacled bear | Chaparri | Zoo Basel | SAMEA3749107 | ERR946788 | Kumar, et al. [3] |
| *Tremarctos ornatus* | Spectacled bear | Nobody | Zoo Basel | SAMEA3749205 | ERR946789 | Kumar, et al. [3] |
| *Ursus maritimus* | Polar bear | PB1 | Spitsbergen, Svalbard | SAMN01057636 | SRR518661, SRR518662 | Miller, et al. [4] |
| *Ursus maritimus* | Polar bear | PB9 | Spitsbergen, Svalbard | SAMN01057666 | SRR518686, SRR518687 | Miller, et al. [4] |
| *Ursus arctos* | Brown bear | GP01 | Glacier National Park, Montana | SAMN02256322 | SRR935609, SRR935616, SRR935617, SRR941811, SRR941814 | Liu, et al. [5] |
| *Ursus americanus* | American black bear | JC012 | Pennsylvania | SAMN02045561 | SRR830685 | Cahill, et al. [6] |
| *Ursus thibetanus* | Asiatic black bear | Anorexica | Zoo Madrid | SAMEA3749106 | ERR946787 | Kumar, et al. [3] |
| *Ursus ursinus* | Sloth bear | Renate | Zoo Liepzig | SAMEA3749105 | ERR946786 | Kumar, et al. [3] |
| *Ursus malayanus* | Sun bear | Anabell | Zoo Munster | SAMEA3749104 | ERR946784 | Kumar, et al. [3] |
| *Ursus malayanus* | Sun bear | Klaus | Zoo Madrid | SAMEA3750870 | ERR946785 | Kumar, et al. [3] |
| *Ailuropoda melanoleuca* | Giant Panda | GP13 | Baoxing, Sichuan | SAMN01040418 | SRR504866 | Zhao, et al. [7] |
| *Ailuropoda melanoleuca* | Giant Panda | GP18 | Beichuan, Sichuan | SAMN01040423 | SRR504871 | Zhao, et al. [7] |

**Table S2:** Details of published and newly sequenced bears used in this study.

*Newly sequenced samples

**Table S3:** Sequencing and mapping statistics for all bear samples analysed in this study. Columns show the number of filtered raw reads used in mapping, the number of successfully mapped reads to the giant panda reference genome, the number of unique reads that mapped after PCR duplicates were removed, the percentage of original reads that mapped, and the percentage of mapped reads that were PCR duplicates.

| Binomial Name | Sample Name | Retained Reads | Raw Mapped Reads | Unique Mapped Reads | Proportion Mapped Reads | Clonality | Coverage (X) |
| --- | --- | --- | --- | --- | --- | --- | --- |
| **Arctodus simus* | ACAD 344 | 496403229 | 279086649 | 202678836 | 0.562217634 | 0.273778102 | 5.92566319 |
| **Arctotherium sp* | ACAD 3599 | 402733914 | 192677824 | 140025115 | 0.478424631 | 0.273268132 | 3.892443033 |
| **Arctodus simus* | ACAD 438 | 800500649 | 33649637 | 23716909 | 0.04203574 | 0.295180837 | 0.519775289 |
| **Arctodus simus* | ACAD 5177 | 475874477 | 5166918 | 4488594 | 0.010857733 | 0.13128213 | 0.121626851 |
| *Tremarctos ornatus* | Chaparri | 312051217 | 268627345 | 256263228 | 0.860843767 | 0.046027023 | 9.199861216 |
| *Tremarctos ornatus* | Nobody | 318173650 | 274620841 | 263660657 | 0.863116229 | 0.039910241 | 9.459313992 |
| *Ursus maritimus* | PB1 | 346660533 | 298508499 | 289412061 | 0.861097444 | 0.030472962 | 11.58217001 |
| *Ursus maritimus* | PB9 | 340345215 | 297785719 | 290239526 | 0.874951978 | 0.025341017 | 11.58504902 |
| *Ursus arctos* | GP01 | 553954695 | 433336259 | 382711352 | 0.782259385 | 0.11682592 | 15.30793958 |
| *Ursus americanus* | JC012 | 194019255 | 160004491 | 153521351 | 0.824683566 | 0.040518488 | 10.34896241 |
| *Ursus thibetanus* | Anorexica | 331174468 | 280801772 | 269428174 | 0.847896801 | 0.040504011 | 9.659247534 |
| *Ursus ursinus* | Renate | 295156761 | 254126145 | 240552726 | 0.860987037 | 0.053412131 | 8.639123446 |
| *Ursus malayanus* | Anabell | 294000644 | 250655873 | 243756105 | 0.852569129 | 0.027526856 | 8.753848913 |
| *Ursus malayanus* | Klaus | 319734539 | 271802375 | 260802737 | 0.850087625 | 0.040469249 | 9.370637783 |
| *Ailuropoda melanoleuca* | GP13 | 137650116 | 119980120 | 111763758 | 0.871631085 | 0.068481028 | 3.765764372 |
| *Ailuropoda melanoleuca* | GP18 | 138649930 | 124714455 | 112803002 | 0.899491655 | 0.095509803 | 4.014613177 |

*Newly sequenced samples

**Table S4:** D-statistics testing for Ursinae hybridisation within short-faced bears (Tremarctinae) using the giant panda as an outgroup. D-statistics (D), standard error, and Z-Score (significant if > |3|) are displayed, with ABBA-BABA counts and the number of SNPs considered in the analysis

| D-statistic: D(H1, H2, H3, Giant Panda) | D | Stderr | Z-score | BABA | ABBA | nSNPs |
| --- | --- | --- | --- | --- | --- | --- |
| D(*T. ornatus* (Nobody), *Arctotherium* sp., *U. maritimus* (PB1)) | 0.1922 | 0.007582 | 25.347 | 10828 | 7336 | 6158376 |
| D(*T. ornatus* (Nobody), *Arctotherium* sp., *U. malayanus* (Klaus)) | 0.2057 | 0.008348 | 24.646 | 10960 | 7219 | 6157958 |
| D(*T. ornatus* (Nobody), *Arctotherium* sp., *U. thibetanus*) | 0.2043 | 0.008319 | 24.555 | 11051 | 7301 | 6171003 |
| D(*T. ornatus* (Nobody), *Arctotherium* sp., *U. malayanus* (Anabell)) | 0.2024 | 0.008262 | 24.499 | 10921 | 7243 | 6158753 |
| D(*T. ornatus* (Nobody), *Arctotherium* sp., *U. ursinus*) | 0.2008 | 0.00824 | 24.367 | 10882 | 7242 | 6163089 |
| D(*T. ornatus* (Nobody), *Arctotherium* sp., *U. maritimus* (PB9)) | 0.1932 | 0.008019 | 24.091 | 10935 | 7394 | 6184441 |
| D(*T. ornatus* (Chaparri), *Arctotherium* sp., *U. malayanus* (Klaus)) | 0.2083 | 0.008725 | 23.875 | 10997 | 7205 | 6159057 |
| D(*T. ornatus* (Chaparri), *Arctotherium* sp., *U. maritimus* (PB1)) | 0.1942 | 0.008137 | 23.86 | 10864 | 7331 | 6158997 |
| D(*T. ornatus* (Chaparri), *Arctotherium* sp., *U. malayanus* (Anabell)) | 0.206 | 0.008636 | 23.851 | 10976 | 7227 | 6159872 |
| D(*T. ornatus* (Nobody), *Arctotherium* sp., *U. arctos*) | 0.2079 | 0.008878 | 23.418 | 11177 | 7329 | 6184195 |
| D(*T. ornatus* (Chaparri), *Arctotherium* sp., *U. thibetanus*) | 0.2052 | 0.008794 | 23.338 | 11072 | 7300 | 6172374 |
| D(*T. ornatus* (Nobody), *Arctotherium* sp., *U. americanus*) | 0.187 | 0.008062 | 23.191 | 10684 | 7318 | 6158065 |
| D(*T. ornatus* (Chaparri), *Arctotherium* sp., *U. ursinus*) | 0.2 | 0.00868 | 23.039 | 10885 | 7256 | 6164301 |
| D(*T. ornatus* (Chaparri), *Arctotherium* sp., *U. americanus*) | 0.188 | 0.008193 | 22.952 | 10727 | 7331 | 6158709 |
| D(*T. ornatus* (Chaparri), *Arctotherium* sp., *U. maritimus* (PB9)) | 0.1962 | 0.008646 | 22.692 | 10978 | 7376 | 6184907 |
| D(*T. ornatus* (Chaparri), *Arctotherium* sp., *U. arctos*) | 0.2074 | 0.009242 | 22.442 | 11185 | 7342 | 6185696 |
| D(*T. ornatus* (Nobody), *A. simus*, *U. arctos*) | 0.2152 | 0.009651 | 22.302 | 11172 | 7214 | 6317724 |
| D(*T. ornatus* (Nobody), *A. simus*, *U. maritimus* (PB1)) | 0.1958 | 0.009013 | 21.728 | 10753 | 7230 | 6290704 |
| D(*T. ornatus* (Nobody), *A. simus*, *U. malayanus* (Klaus)) | 0.2065 | 0.009668 | 21.36 | 10878 | 7154 | 6290343 |
| D(*T. ornatus* (Chaparri), *A. simus*, *U. arctos*) | 0.2131 | 0.010012 | 21.281 | 11191 | 7260 | 6319318 |
| D(*T. ornatus* (Nobody), *A. simus*, *U. thibetanus*) | 0.2091 | 0.009899 | 21.126 | 11026 | 7212 | 6302970 |
| D(*T. ornatus* (Nobody), *A. simus*, *U. maritimus* (PB9)) | 0.2003 | 0.009484 | 21.125 | 10898 | 7259 | 6317216 |
| D(*T. ornatus* (Chaparri), *A. simus*, *U. thibetanus*) | 0.2104 | 0.010212 | 20.604 | 11076 | 7225 | 6304295 |
| D(*T. ornatus* (Nobody), *A. simus*, *U. americanus*) | 0.1879 | 0.00916 | 20.512 | 10580 | 7233 | 6292779 |
| D(*T. ornatus* (Chaparri), *A. simus*, *U. malayanus* (Klaus)) | 0.2075 | 0.010129 | 20.487 | 10909 | 7159 | 6291437 |
| D(*T. ornatus* (Nobody), *A. simus*, *U. malayanus* (Anabell)) | 0.2029 | 0.009971 | 20.347 | 10862 | 7198 | 6290304 |
| D(*T. ornatus* (Chaparri), *A. simus*, *U. maritimus* (PB9)) | 0.1998 | 0.009847 | 20.287 | 10942 | 7298 | 6317882 |
| D(*T. ornatus* (Nobody), *A. simus*, *U. ursinus*) | 0.206 | 0.010293 | 20.014 | 10868 | 7154 | 6294596 |
| D(*T. ornatus* (Chaparri), *A. simus*, *U. maritimus* (PB1)) | 0.196 | 0.01 | 19.604 | 10789 | 7252 | 6291256 |
| D(*T. ornatus* (Chaparri), *A. simus*, *U. malayanus* (Anabell)) | 0.2042 | 0.010502 | 19.444 | 10879 | 7189 | 6291443 |
| D(*T. ornatus* (Chaparri), *A. simus*, *U. americanus*) | 0.188 | 0.009687 | 19.403 | 10609 | 7252 | 6293439 |
| D(*T. ornatus* (Chaparri), *A. simus*, *U. ursinus*) | 0.2036 | 0.010578 | 19.246 | 10855 | 7182 | 6295812 |

**Table S5:** D-statistics testing for Tremarctinae hybridisation within Ursinae using the giant panda as an outgroup. D-statistics (D), standard error, and Z-Score (significant if > |3|) are displayed, with ABBA-BABA counts and the number of SNPs considered in the analysis

| D-statistic: D(H1, H2, H3, Giant Panda) | D | Stderr | Z-score | BABA | ABBA | nSNPs |
| --- | --- | --- | --- | --- | --- | --- |
| D(*U. arctos*, *U. americanus*, *T. ornatus* (Nobody)) | 0.038 | 0.006774 | 5.615 | 10688 | 9904 | 6585424 |
| D(*U. arctos*, *U. maritimus* (PB1), *T. ornatus* (Nobody)) | 0.0664 | 0.012707 | 5.222 | 6824 | 5974 | 6612645 |
| D(*U. arctos*, *U. maritimus* (PB9), *T. ornatus* (Nobody)) | 0.0639 | 0.012328 | 5.181 | 6869 | 6043 | 6646165 |
| D(*U. arctos*, *U. maritimus* (PB1), *T. ornatus* (Chaparri)) | 0.0639 | 0.012684 | 5.041 | 6867 | 6041 | 6614829 |
| D(*U. arctos*, *U. maritimus* (PB9), *T. ornatus* (Chaparri)) | 0.061 | 0.012929 | 4.717 | 6847 | 6059 | 6648311 |
| D(*U. arctos*, *U. americanus*, *T. ornatus* (Chaparri)) | 0.0309 | 0.006752 | 4.577 | 10651 | 10011 | 6587600 |
| D(*U. thibetanus*, *U. maritimus* (PB1), *T. ornatus* (Chaparri)) | 0.0377 | 0.009127 | 4.136 | 12758 | 11829 | 6580987 |
| D(*U. thibetanus*, *U. maritimus* (PB1), *T. ornatus* (Nobody)) | 0.0378 | 0.009426 | 4.013 | 12691 | 11764 | 6579114 |
| D(*U. ursinus*, *U. maritimus* (PB1), *T. ornatus* (Chaparri)) | 0.0252 | 0.006652 | 3.794 | 14671 | 13948 | 6566596 |
| D(*U. ursinus*, *U. maritimus* (PB1), *T. ornatus* (Nobody)) | 0.0253 | 0.006734 | 3.761 | 14617 | 13894 | 6565035 |
| D(*U. thibetanus*, *U. maritimus* (PB9), *T. ornatus* (Chaparri)) | 0.0318 | 0.009213 | 3.447 | 12689 | 11907 | 6614426 |
| D(*U. thibetanus*, *U. americanus*, *T. ornatus* (Nobody)) | 0.0315 | 0.009223 | 3.412 | 12659 | 11886 | 6562354 |
| D(*U. thibetanus*, *U. maritimus* (PB9), *T. ornatus* (Nobody)) | 0.0309 | 0.009438 | 3.279 | 12628 | 11869 | 6612456 |
| D(*U. ursinus*, *U. maritimus* (PB9), *T. ornatus* (Nobody)) | 0.0216 | 0.006665 | 3.242 | 14605 | 13986 | 6598187 |
| D(*U. thibetanus*, *U. americanus*, *T. ornatus* (Chaparri)) | 0.0287 | 0.008944 | 3.214 | 12632 | 11925 | 6564284 |
| D(*U. ursinus*, *U. maritimus* (PB9), *T. ornatus* (Chaparri)) | 0.0211 | 0.006599 | 3.199 | 14624 | 14018 | 6599795 |
| D(*U. ursinus*, *U. americanus*, *T. ornatus* (Nobody)) | 0.02 | 0.00684 | 2.919 | 14280 | 13721 | 6549440 |
| D(*U. thibetanus*, *U. malayanus* (Anabell), *Arctotherium* sp.) | 0.0219 | 0.007818 | 2.807 | 9969 | 9540 | 6169185 |
| D(*U. thibetanus*, *U. malayanus* (Klaus), *Arctotherium* sp.) | 0.0208 | 0.007595 | 2.734 | 9997 | 9590 | 6166665 |
| D(*U. malayanus* (Anabell), *U. maritimus* (PB1), *T. ornatus* (Chaparri)) | 0.0182 | 0.006915 | 2.639 | 14490 | 13970 | 6562190 |
| D(*U. thibetanus*, *U. malayanus* (Klaus), *A. simus*) | 0.0219 | 0.008346 | 2.625 | 10302 | 9859 | 6298833 |
| D(*U. ursinus*, *U. americanus*, *T. ornatus* (Chaparri)) | 0.0173 | 0.006651 | 2.597 | 14286 | 13801 | 6551084 |
| D(*U. thibetanus*, *U. malayanus* (Anabell), *A. simus*) | 0.0219 | 0.008617 | 2.544 | 10267 | 9826 | 6300444 |
| D(*U. malayanus* (Anabell), *U. maritimus* (PB1), *T. ornatus* (Nobody)) | 0.0171 | 0.006838 | 2.496 | 14405 | 13922 | 6560582 |
| D(*U. thibetanus*, *U. malayanus* (Anabell), *T. ornatus* (Nobody)) | 0.0209 | 0.008549 | 2.439 | 11846 | 11361 | 6621261 |
| D(*U. malayanus* (Klaus), *U. maritimus* (PB1), *T. ornatus* (Nobody)) | 0.0179 | 0.007362 | 2.426 | 14489 | 13981 | 6559995 |
| D(*U. arctos*, *U. malayanus* (Anabell), *T. ornatus* (Nobody)) | 0.0159 | 0.006604 | 2.404 | 14306 | 13858 | 6630767 |
| D(*U. malayanus* (Klaus), *U. maritimus* (PB1), *T. ornatus* (Chaparri)) | 0.017 | 0.007132 | 2.386 | 14515 | 14030 | 6561717 |
| D(*U. thibetanus*, *U. malayanus* (Anabell), *T. ornatus* (Chaparri)) | 0.0198 | 0.008296 | 2.382 | 11879 | 11418 | 6623482 |
| D(*U. ursinus*, *U. malayanus* (Anabell), *Arctotherium* sp.) | 0.0172 | 0.007243 | 2.373 | 9061 | 8754 | 6162684 |
| D(*U. malayanus* (Klaus), *U. americanus*, *T. ornatus* (Nobody)) | 0.0135 | 0.006132 | 2.197 | 14115 | 13740 | 6545186 |
| D(*U. americanus*, *U. malayanus* (Anabell), *Arctotherium* sp.) | 0.015 | 0.006894 | 2.17 | 12125 | 11766 | 6146174 |
| D(*U. thibetanus*, *U. malayanus* (Klaus), *T. ornatus* (Chaparri)) | 0.0183 | 0.008645 | 2.119 | 11898 | 11469 | 6621804 |
| D(*U. americanus*, *U. malayanus* (Klaus), *Arctotherium* sp.) | 0.014 | 0.006638 | 2.109 | 12111 | 11776 | 6146598 |
| D(*U. ursinus*, *U. malayanus* (Klaus), *Arctotherium* sp.) | 0.0161 | 0.007726 | 2.084 | 9078 | 8790 | 6160076 |
| D(*U. arctos*, *U. malayanus* (Anabell), *T. ornatus* (Chaparri)) | 0.0139 | 0.006701 | 2.075 | 14310 | 13917 | 6633179 |
| D(*U. arctos*, *U. malayanus* (Klaus), *T. ornatus* (Chaparri)) | 0.0131 | 0.006487 | 2.017 | 14293 | 13923 | 6632784 |
| D(*U. arctos*, *U. malayanus* (Klaus), *T. ornatus* (Nobody)) | 0.013 | 0.006487 | 1.999 | 14267 | 13901 | 6630244 |
| D(*U. thibetanus*, *U. malayanus* (Klaus), *T. ornatus* (Nobody)) | 0.0182 | 0.009231 | 1.972 | 11827 | 11403 | 6619492 |
| D(*U. americanus*, *U. malayanus* (Anabell), *A. simus*) | 0.0125 | 0.006756 | 1.854 | 12469 | 12159 | 6279339 |
| D(*U. malayanus* (Anabell), *U. maritimus* (PB9), *T. ornatus* (Chaparri)) | 0.0129 | 0.006962 | 1.852 | 14438 | 14070 | 6595307 |
| D(*U. malayanus* (Klaus), *U. maritimus* (PB9), *T. ornatus* (Nobody)) | 0.0131 | 0.007208 | 1.812 | 14448 | 14075 | 6593005 |
| D(*U. malayanus* (Anabell), *U. maritimus* (PB9), *T. ornatus* (Nobody)) | 0.0123 | 0.006926 | 1.775 | 14403 | 14052 | 6593765 |
| D(*U. malayanus* (Klaus), *U. americanus*, *T. ornatus* (Chaparri)) | 0.0107 | 0.00619 | 1.734 | 14152 | 13852 | 6546765 |
| D(*U. malayanus* (Anabell), *U. americanus*, *T. ornatus* (Nobody)) | 0.0106 | 0.006177 | 1.712 | 14113 | 13817 | 6545768 |
| D(*U. americanus*, *U. malayanus* (Klaus), *A. simus*) | 0.0115 | 0.006802 | 1.687 | 12471 | 12187 | 6280782 |
| D(*U. malayanus* (Klaus), *U. maritimus* (PB9), *T. ornatus* (Chaparri)) | 0.0108 | 0.007032 | 1.538 | 14429 | 14120 | 6594752 |
| D(*U. thibetanus*, *U. maritimus* (PB1), *Arctotherium* sp.) | 0.0124 | 0.00808 | 1.536 | 10388 | 10133 | 6160785 |
| D(*U. ursinus*, *U. malayanus* (Anabell), *A. simus*) | 0.012 | 0.008027 | 1.496 | 9221 | 9002 | 6293600 |
| D(*U. ursinus*, *U. malayanus* (Klaus), *A. simus*) | 0.0119 | 0.008029 | 1.484 | 9238 | 9020 | 6291880 |
| D(*U. ursinus*, *U. malayanus* (Klaus), *T. ornatus* (Chaparri)) | 0.0112 | 0.007567 | 1.481 | 10795 | 10556 | 6611944 |
| D(*U. ursinus*, *U. malayanus* (Anabell), *T. ornatus* (Nobody)) | 0.0104 | 0.007157 | 1.45 | 10715 | 10495 | 6611714 |
| D(*U. malayanus* (Anabell), *U. americanus*, *T. ornatus* (Chaparri)) | 0.0092 | 0.006374 | 1.447 | 14136 | 13877 | 6547286 |
| D(*U. maritimus* (PB9), *U. malayanus* (Anabell), *A. simus*) | 0.0097 | 0.006692 | 1.442 | 12513 | 12274 | 6304388 |
| D(*U. arctos*, *U. malayanus* (Klaus), *A. simus*) | 0.0084 | 0.005941 | 1.422 | 12439 | 12230 | 6306227 |
| D(*U. maritimus* (PB9), *U. malayanus* (Anabell), *Arctotherium* sp.) | 0.0095 | 0.00683 | 1.396 | 12168 | 11938 | 6172713 |
| D(*U. americanus*, *U. maritimus* (PB1), *Arctotherium* sp.) | 0.0097 | 0.006981 | 1.383 | 8712 | 8545 | 6163675 |
| D(*U. maritimus* (PB9), *U. malayanus* (Klaus), *A. simus*) | 0.0092 | 0.006852 | 1.348 | 12505 | 12275 | 6305556 |
| D(*U. arctos*, *U. malayanus* (Anabell), *A. simus*) | 0.0081 | 0.006004 | 1.344 | 12435 | 12235 | 6306804 |
| D(*U. maritimus* (PB9), *U. malayanus* (Klaus), *Arctotherium* sp.) | 0.0088 | 0.006675 | 1.324 | 12157 | 11944 | 6172885 |
| D(*U. maritimus* (PB1), *U. maritimus* (PB9), *Arctotherium* sp.) | 0.0326 | 0.025583 | 1.276 | 816 | 764 | 6217830 |
| D(*U. ursinus*, *U. malayanus* (Klaus), *T. ornatus* (Nobody)) | 0.0104 | 0.008174 | 1.272 | 10737 | 10515 | 6609836 |
| D(*U. arctos*, *U. malayanus* (Anabell), *Arctotherium* sp.) | 0.0093 | 0.007391 | 1.253 | 12077 | 11856 | 6174614 |
| D(*U. maritimus* (PB1), *U. maritimus* (PB9), *A. simus*) | 0.0361 | 0.029276 | 1.234 | 850 | 790 | 6352025 |
| D(*U. arctos*, *U. malayanus* (Klaus), *Arctotherium* sp.) | 0.0089 | 0.007464 | 1.189 | 12073 | 11860 | 6173076 |
| D(*U. ursinus*, *U. malayanus* (Anabell), *T. ornatus* (Chaparri)) | 0.009 | 0.007655 | 1.174 | 10700 | 10509 | 6613671 |
| D(*U. thibetanus*, *U. ursinus*, *T. ornatus* (Chaparri)) | 0.0087 | 0.007445 | 1.169 | 12094 | 11885 | 6627699 |
| D(*U. thibetanus*, *U. maritimus* (PB9), *Arctotherium* sp.) | 0.0089 | 0.00805 | 1.108 | 10410 | 10225 | 6186620 |
| D(*U. thibetanus*, *U. arctos*, *A. simus*) | 0.009 | 0.008333 | 1.075 | 10491 | 10303 | 6323067 |
| D(*U. thibetanus*, *U. arctos*, *Arctotherium* sp.) | 0.009 | 0.008403 | 1.075 | 10232 | 10048 | 6190143 |
| D(*U. thibetanus*, *U. maritimus* (PB1), *A. simus*) | 0.0093 | 0.008807 | 1.051 | 10576 | 10381 | 6292413 |
| D(*U. thibetanus*, *U. ursinus*, *A. simus*) | 0.0101 | 0.009698 | 1.044 | 10471 | 10260 | 6303292 |
| D(*U. thibetanus*, *U. ursinus*, *Arctotherium* sp.) | 0.0077 | 0.007388 | 1.043 | 10153 | 9997 | 6172199 |
| D(*U. thibetanus*, *U. ursinus*, *T. ornatus* (Nobody)) | 0.0081 | 0.007806 | 1.039 | 12052 | 11858 | 6625682 |
| D(*U. thibetanus*, *U. maritimus* (PB9), *A. simus*) | 0.0083 | 0.008649 | 0.956 | 10587 | 10412 | 6318797 |
| D(*U. maritimus* (PB1), *U. malayanus* (Anabell), *A. simus*) | 0.0063 | 0.006685 | 0.946 | 12449 | 12292 | 6277891 |
| D(*U. ursinus*, *U. maritimus* (PB1), *Arctotherium* sp.) | 0.0061 | 0.006559 | 0.924 | 12233 | 12086 | 6149992 |
| D(*U. maritimus* (PB1), *U. malayanus* (Anabell), *Arctotherium* sp.) | 0.0059 | 0.006736 | 0.87 | 12080 | 11939 | 6147000 |
| D(*U. americanus*, *U. maritimus* (PB1), *A. simus*) | 0.0065 | 0.007941 | 0.816 | 8965 | 8849 | 6298576 |
| D(*U. thibetanus*, *U. americanus*, *Arctotherium* sp.) | 0.0055 | 0.007927 | 0.689 | 10461 | 10347 | 6158742 |
| D(*U. americanus*, *U. maritimus* (PB9), *Arctotherium* sp.) | 0.005 | 0.007409 | 0.673 | 8722 | 8635 | 6189156 |
| D(*U. americanus*, *U. maritimus* (PB1), *T. ornatus* (Chaparri)) | 0.0047 | 0.007502 | 0.626 | 10245 | 10149 | 6548802 |
| D(*U. ursinus*, *U. maritimus* (PB9), *Arctotherium* sp.) | 0.0041 | 0.006544 | 0.623 | 12225 | 12126 | 6175763 |
| D(*U. maritimus* (PB1), *U. malayanus* (Klaus), *A. simus*) | 0.0044 | 0.007094 | 0.615 | 12432 | 12323 | 6279261 |
| D(*U. ursinus*, *U. arctos*, *Arctotherium* sp.) | 0.0044 | 0.007176 | 0.613 | 12135 | 12028 | 6178377 |
| D(*U. arctos*, *U. ursinus*, *T. ornatus* (Nobody)) | 0.0045 | 0.007454 | 0.609 | 14264 | 14135 | 6635967 |
| D(*U. thibetanus*, *U. americanus*, *A. simus*) | 0.005 | 0.008311 | 0.603 | 10769 | 10662 | 6292658 |
| D(*U. maritimus* (PB1), *U. malayanus* (Klaus), *Arctotherium* sp.) | 0.004 | 0.006962 | 0.576 | 12052 | 11954 | 6147130 |
| D(*U. americanus*, *U. arctos*, *Arctotherium* sp.) | 0.0041 | 0.007475 | 0.546 | 8699 | 8628 | 6176832 |
| D(*U. ursinus*, *U. maritimus* (PB1), *A. simus*) | 0.0036 | 0.006714 | 0.542 | 12472 | 12382 | 6281231 |
| D(*U. arctos*, *U. ursinus*, *T. ornatus* (Chaparri)) | 0.0037 | 0.00755 | 0.495 | 14265 | 14158 | 6638300 |
| D(*U. malayanus* (Anabell), *U. malayanus* (Klaus), *T. ornatus* (Chaparri)) | 0.0072 | 0.015624 | 0.46 | 2495 | 2459 | 6638489 |
| D(*U. arctos*, *U. maritimus* (PB9), *A. simus*) | 0.0049 | 0.011036 | 0.441 | 5348 | 5296 | 6347717 |
| D(*U. americanus*, *U. arctos*, *A. simus*) | 0.0033 | 0.007697 | 0.431 | 8976 | 8917 | 6312319 |
| D(*U. arctos*, *U. maritimus* (PB1), *A. simus*) | 0.0046 | 0.01109 | 0.412 | 5336 | 5287 | 6321140 |
| D(*U. americanus*, *U. maritimus* (PB9), *A. simus*) | 0.003 | 0.008203 | 0.36 | 8960 | 8906 | 6324725 |
| D(*U. ursinus*, *U. arctos*, *A. simus*) | 0.0023 | 0.006705 | 0.349 | 12408 | 12350 | 6310873 |
| D(*U. arctos*, *U. maritimus* (PB1), *Arctotherium* sp.) | 0.0037 | 0.01124 | 0.325 | 5117 | 5080 | 6187644 |
| D(*U. ursinus*, *U. maritimus* (PB9), *A. simus*) | 0.002 | 0.006596 | 0.308 | 12441 | 12390 | 6307573 |
| D(*U. americanus*, *U. ursinus*, *Arctotherium* sp.) | 0.0016 | 0.00651 | 0.24 | 12034 | 11997 | 6148976 |
| D(*U. americanus*, *U. maritimus* (PB1), *T. ornatus* (Nobody)) | 0.0017 | 0.007694 | 0.215 | 10165 | 10131 | 6547890 |
| D(*U. thibetanus*, *U. arctos*, *T. ornatus* (Chaparri)) | 0.0021 | 0.010081 | 0.206 | 12079 | 12028 | 6654575 |
| D(*U. malayanus* (Anabell), *U. malayanus* (Klaus), *A. simus*) | 0.0028 | 0.015051 | 0.189 | 1899 | 1888 | 6316094 |
| D(*U. maritimus* (PB1), *U. maritimus* (PB9), *T. ornatus* (Nobody)) | 0.0044 | 0.023285 | 0.188 | 1242 | 1231 | 6631992 |
| D(*U. malayanus* (Anabell), *U. malayanus* (Klaus), *Arctotherium* sp.) | 0.0031 | 0.01673 | 0.187 | 1854 | 1842 | 6183689 |
| D(*U. arctos*, *U. maritimus* (PB9), *Arctotherium* sp.) | 0.0021 | 0.011285 | 0.187 | 5150 | 5128 | 6213415 |
| D(*U. americanus*, *U. ursinus*, *A. simus*) | 0.0011 | 0.007109 | 0.159 | 12368 | 12339 | 6282149 |
| D(*U. arctos*, *U. thibetanus*, *T. ornatus* (Nobody)) | 0.0016 | 0.010223 | 0.154 | 12068 | 12031 | 6651971 |
| D(*U. maritimus* (PB9), *U. americanus*, *T. ornatus* (Nobody)) | 0.001 | 0.008065 | 0.13 | 10221 | 10201 | 6577928 |
| D(*U. maritimus* (PB9), *U. americanus*, *T. ornatus* (Chaparri)) | 0.001 | 0.00799 | 0.126 | 10262 | 10242 | 6578875 |
| D(*U. maritimus* (PB9), *U. maritimus* (PB1), *T. ornatus* (Chaparri)) | 0.0013 | 0.0221 | 0.058 | 1284 | 1281 | 6633124 |
| D(*U. malayanus* (Klaus), *U. malayanus* (Anabell), *T. ornatus* (Nobody)) | 0.0001 | 0.014705 | 0.006 | 2451 | 2451 | 6636253 |
